# iDamIDseq and iDEAR: An improved method and a computational pipeline to profile chromatin-binding proteins of developing organisms

**DOI:** 10.1101/062208

**Authors:** Jose Arturo Gutierrez-Triana, Juan L. Mateo, David Ibberson, Joachim Wittbrodt

## Abstract

DNA adenine methyltransferase identification (DamID) has emerged as an alternative for profiling protein-DNA interactions, however critical issues in the method limit its applicability. Here we present iDamIDseq, a protocol that improves specificity and robustness making its use compatible with developing organisms. In addition, we present the analysis tool iDEAR (iDamIDseq Enrichment Analysis with R) to determine protein-DNA interactions genome wide. The combination of both allows establishing highly reliable transcription factor profiles, even in transient assays. For tissue specific expression we improved the Dam coding sequence to overcome predominant aberrant splicing of Dam fusions we discovered with the commonly used sequence.

## Introduction

Animal development is the result of an exquisite orchestration of changes in gene expression in time and space. Transcription factors (TF) and other chromatin-associated proteins are fundamental elements in these processes and the search for their targets and they logic of regulation in the genome is matter of deep investigation. Chromatin immunopreicpitation (ChIP) and DNA adenine methyltransferase identification (DamID) are the prevailing methods to profile transcription factor binding regions in the genome, reviewed in (Aughey and Southall, 2016; Furey, 2012). ChIP relies on antibody-based capture of protein-DNA complexes on crosslinked and sheared chromatin. Although the protocol is robust, it depends on highly specific precipitating antibodies. In particular, crossreacting antibodies may immunoprecipitate more than one TF simultaneously in a ChIP experiment. DamID offers a suitable solution to these problems. In DamID the fusion of a TF to the bacterial gene DNA adenine methyltransferase, *Dam*, allows for a restricted methylation of adenine residues of the GATC target sequences near the TF binding sites. These regions can then be selected and enriched by digesting the DNA with the restriction enzyme DpnI and linker-mediated PCR (LM-PCR). The PCR products are then hybridized to microarrays or used directly for deep sequencing. DamID is a technique that requires relatively low input material, less processing time and it is cost-effective. DamID has been used successfully in different model organisms such as *Drosophila melanogaster* (Southall et al., 2013), *Caenorhabditis elegans* (Schuster et al., 2010), *Arabidopsis thaliana* (Germann et al., 2006) and mammalian cell culture (Vogel et al., 2007). However, the current protocols require tight control to ensure low expression levels of the proteins fused to the *E.coli* Dam methylase. In addition, in the process of implementing DamID to developing medaka embryos, we faced serious problems such as lack of any DamID product and nonspecific amplification (DpnI-independent amplification).

To overcome these drawbacks and allow wide and immediate application of the approach we have implemented iDamIDseq with a series of improvements to the original protocol resulting in a robust and easily applicable method. The approach presented here now allows transcription factor profiling even in transient applications. We complement our experimental improvements with iDEAR (iDamID Enrichment Analysis with R), an associated analysis pipeline to swiftly establish highly reliable transcription factor profiles.

## Results and Discussion

The iDamIDseq protocol comprises modifications to critical steps of the original technique described below (Fig.1).

Since the E. coli Dam displays activity in near-cognate sequences (Horton et al., 2005), we used the mutant version DamL122A (herein referred as Dam) to increase the specificity of methylation on GATC sites.

Chimeric fusion proteins may compromise function due to steric hindrance (Arai et al., 2001). Therefore, we included a flexible linker between the *Dam* protein and the transcription factor, and tested different orientations. We observed that the methylation defect of Dam-deficient bacteria could be rescued differentially by the *Dam* fusions depending on the orientation of the fused protein (amino or carboxy terminal) and the lack or presence of the flexylinker (Supplementary Fig.1 and data not shown), these results may explain inefficient DamID amplification in some cases. Noteworthy, medaka embryos injected with mRNA of a fusion of the *Dam* gene and GFP are able to develop properly as assessed at 34 hours post fertilization (Fig.1B,C).

**Figure 1:**
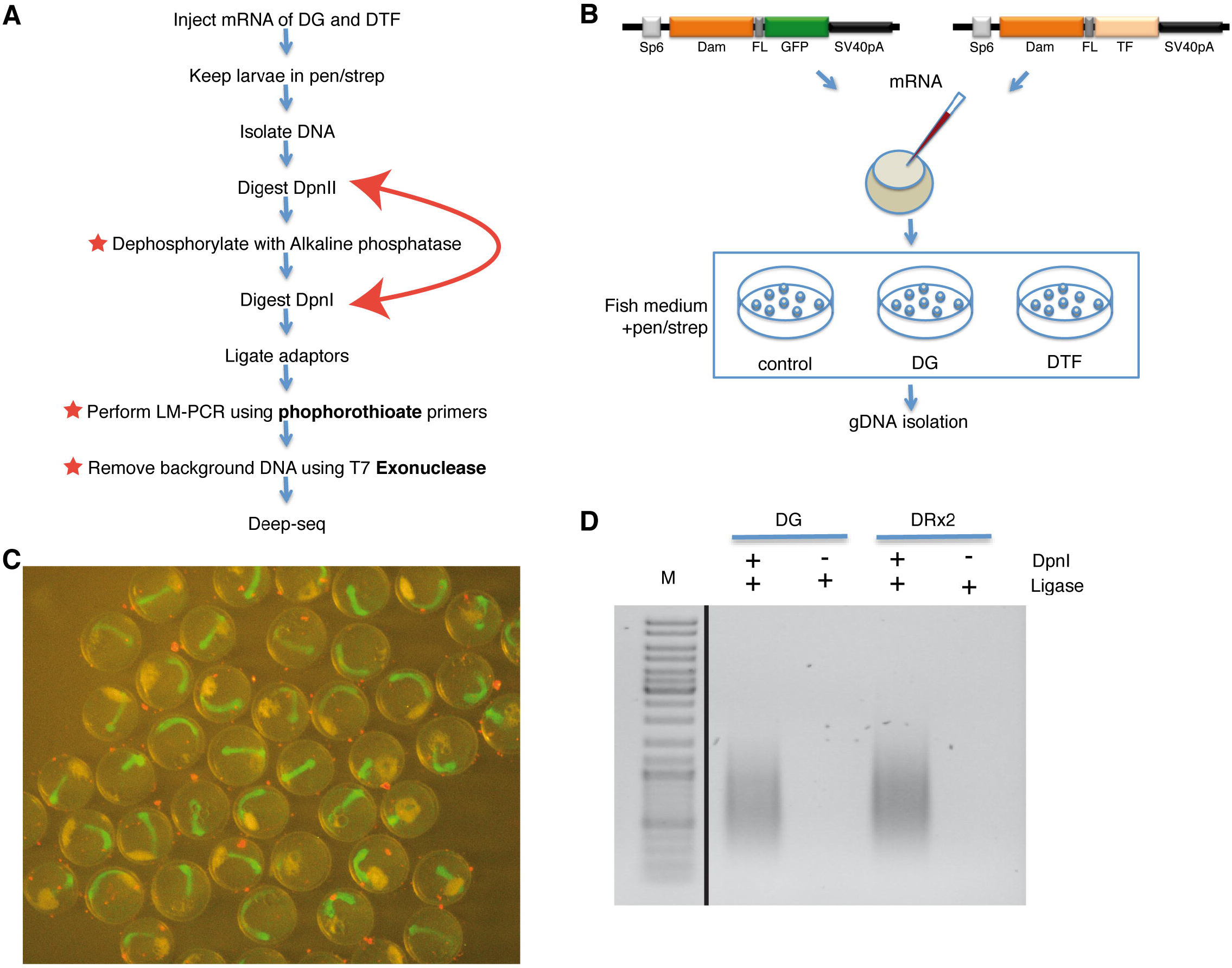
iDamlDseq to robustly analyze protein-DNA interactions. A) Overview of the iDamlDseq procedure. Critical modifications are highlighted in red. B) Delivery of Dam fusion proteins. In vitro synthesized mRNA encoding Dam_GFP (DG) or Dam_TF (DTF) were injected into 1 cell stage medaka zygotes. C) Dam_GFP expression in medaka embryos (stage 22) did not affect embryonic development. D) iDamID amplification at 25 cycles from embryos injected with DG or DRx2 mRNA. Amplified DNA fragments were observed only in samples treated with DpnI. The black line marks intervening lanes removed.

Since we repeatedly obtained linker-mediated amplification independent of DpnI (data not shown), we reasoned that this problem is due to the ligation of the adaptors to free phosphorylated 5´ends resulting from the original genomic DNA preparation and not from the DpnI digestion. We dramatically enhanced the specificity of the adaptor ligation by switching the order of the DpnI and DpnII digestions and the addition of an alkaline phosphatase step. First, we reduced size-complexity by digesting the DNA with DpnII, which cuts GATC sites but is sensitive to adenine methylation. Then, we treated these fragments with alkaline phosphatase and next proceeded with the digestion using DpnI, which only cuts methylated adenine GATC sites. LM-PCR amplification products were obtained only in samples treated with DpnI (Fig.1D).

In order to prepare the sample for deep sequencing, any contaminant genomic DNA must be removed. Therefore, we performed LM-PCR using primers protected with phosphorothioate modifications and next we treated the samples with T7 exonuclease.

As a proof of concept, we applied iDamIDseq to medaka (Oryzias latipes) to reveal a chromatin binding profile of the transcription factor Rx2, the homolog of the mammalian Rax homeodomain proteins involved in retina development We injected mRNA coding for a nuclear localized Dam_GFP or Dam_Rx2 into one cell-stage medaka zygotes and allowed them to develop to Stage 22 (Iwamatsu, 2004) (34 hpf at 28°C, Fig.1B). We extracted the gDNA and processed the sample as described above with two biological replicates per condition. The correlation of read coverage over the genome is very high between replicates and quite distinct between Rx2 and GFP showing the robustness of this method (Fig.2A). We developed an R package, named iDEAR (iDamID Enrichment Analysis with R, available at https://bitbucket.org/juanlmateo/idear), to facilitate the straightforward analysis of differential methylation regions (see Methods). Using iDEAR, we were able to identify 7948 Rx2 target regions. Strikingly, we also identified 6255 regions with a significant depletion of signal in the Rx2 samples with respect to GFP. The average length of Rx2 enriched regions is 916bp while for the Rx2 depleted regions is lOllbp (median 731 and 830 respectively), indicating methylation of discrete and specific regions in the genome. The presence of the RAX binding motif correlated well with the enrichment level (Fig.2B), with over 70% of the top Rx2 target regions containing this motif. This suggests that these regions are *bona fide* Rx2 binding sites. Conversely, only 24% of the regions depleted of Rx2 contain this motif on average. This is clearly below the expected frequency of the homeodomain binding motif in random sequences, indicating a counter-selection against the Rx2 binding motif in these regions.

**Figure 2:**
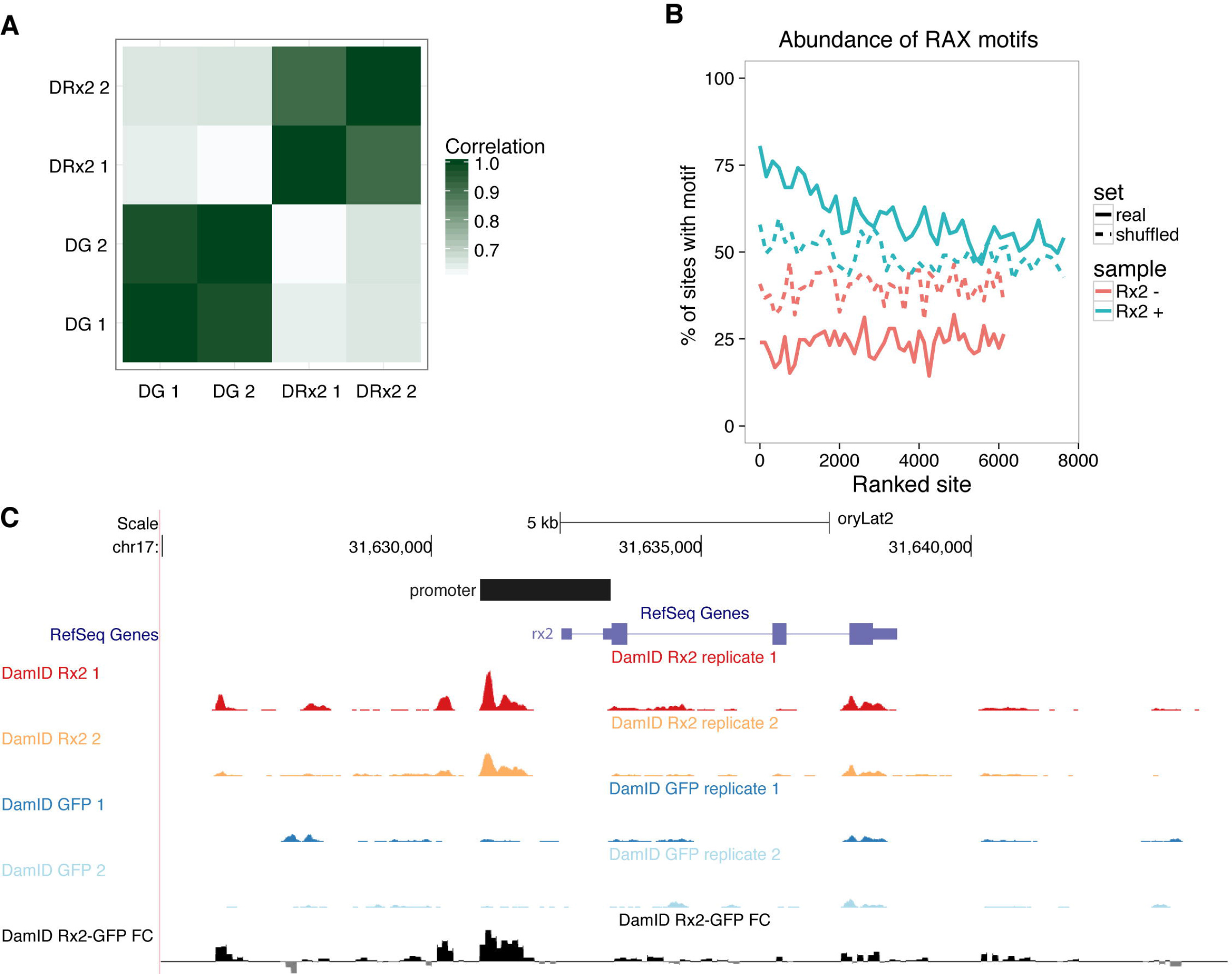
Analysis of iDamlDseq on Rx2. A) Samples showed high correlation between replicates and low correlation between DRx2 and DG based on the genome wide read coverage. B) Representation of the abundance of the RAX binding motif with respect to the level of significance of the identified regions. In blue and red are the Rx2 enriched and depleted regions respectively, dashed lines represent the background established by random shuffling of the same sequence. C) Representation of the iDamlDseq signal around the *Rx2* locus showing the normalized read coverage for each one of the replicates of Rx2 and GFP fusions and the log2 fold change of the average signal per replica (track in black at the bottom). The black bar over the RefSeq Genes track represents the previously characterized Rx2 promoter. The area of maximum Rx2 enrichment is included in the 5’ end of this promoter.

Inspection of regions of Rx2 enrichment revealed many known players involved in retinal development, such as Six3 (Loosli et al., 1998), Otx2 (Zuber et al., 2003), Pax6 (Loosli et al., 1998) and Sox2 (Reinhardt et al., 2015). Interestingly, we also found enrichment of Rx2 on its own proximal upstream locus (Fig.2C). Part of this region overlaps with a previously characterized regulatory sequence labeled as the Rx2 promoter (Reinhardt et al., 2015), suggesting that Rx2 acts in a self-regulatory feedback loop.

Endogenous promoters can be used to generate transcription factor binding profiles in a tissue specific manner. We cloned Dam_GFP after different promoters (ubiquitin (Mosimann et al., 2011), heat shock (Blechinger et al., 2002) and Rx2 (Reinhardt et al., 2015)) in plasmids carrying transgenesis markers such as Cmlc2:GFP or RFP. Surprisingly, none of the Dam_GFP constructs showed GFP expression, whereas the unfused GFP construct did (Fig.3A and data not shown). Since the Dam_GFP can be translated efficiently (see Fig.1C), we suspected of problems at the transcriptional/splicing level. RT_PCR of samples from the different Ubiquitin-driven constructs revealed the aberrant splicing of the *Dam* gene out of the final transcript (Fig.3A–B; Supplementary Fig.2). A customized optimization of the *Dam* gene (*oDam*) removed the cryptic splicing regulatory sites and restored the expression of the GFP in the larvae (Fig.3C; Supplementary Fig.3).

**Figure 3:**
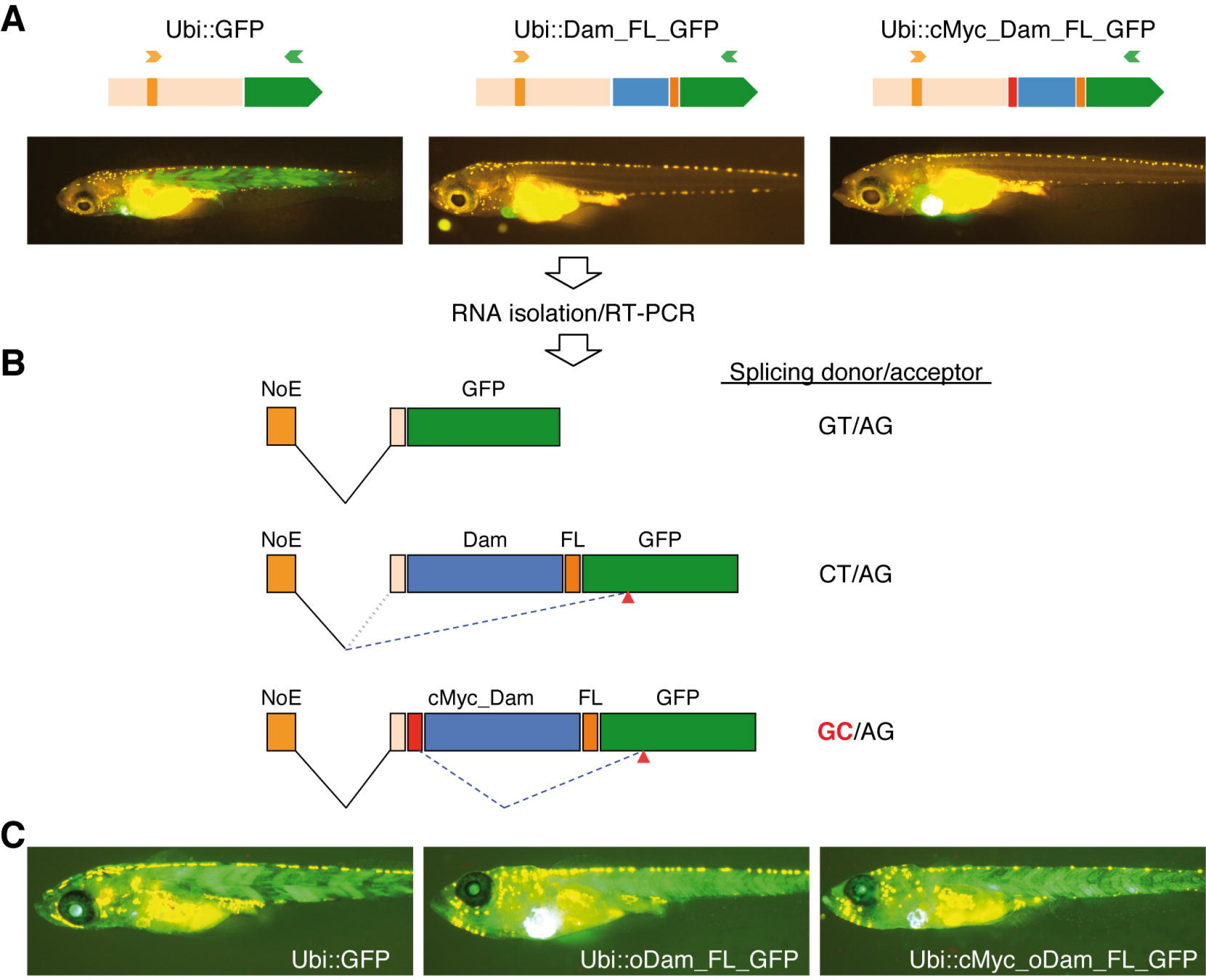
Optimization of the bacterial Dam gene is necessary for proper expression of Dam fusion proteins, avoiding aberrant splicing. A) Plasmids containing GFP, DG and cMyc_DG cassettes driven by the 3.5Kb Ubiquitin promoter (Ubi) were co-injected with Tol2 transposase into one cell stage zygotes. Successfully injected larvae expressing EGFP in the heart were selected for further studies. Note that only Ubi::GFP expressed ubiquitously in the body of the larvae. B) RNA was isolated from pools of larvae from the experimental groups. RT_PCR was performed using a forward primer annealing in the noncoding exon included in Ubiquitin promoter (NoE) and the reverse primer in the body of the GFP coding sequence. Proper splicing occurs between NoE and GFP in the Ubi::GFP larvae. In Ubi::Dam_mGFP larvae, incorrect splicing occurs between the NoE and a cryptic acceptor site in the GFP coding region (red arrowhead). In the Ubi::cMyc_Dam_GFP, NoE is spliced to the proper acceptor upstream of the cMyc sequence, but it is miss-spliced after the cMyc sequence, using a cryptic donor site, to the same cryptic acceptor sequence in the mGFP as for Ubi::Dam_GFP (see also supplementary Fig.3). The prokaryotic Dam ORF carries a strong splicing potentiator recognized in the eukaryotic context C) Optimization of the Dam ORF removed this potential, facilitating proper expression of the fusion proteins.

In conclusion, iDamlDseq generates reliable and robust transcription factor profiles when used in developing medaka embryos. Since the *Dam* is a prokaryotic gene, care must be taken to not only optimize codon usage but also eliminate any cryptic splicing sites to achieve proper expression of the fusion protein in different tissues of the fish. This technique can be extended to determine the binding profile of transcription factors of animals transiently via injection in early stages of either RNA or DNA plasmids. Furthermore, these improvements will allow implementing DamID in any organism that is amenable for transgenesis allowing a tissue and time specific profile.

## Materials and methods

### Fish maintenance

Medaka (*Oryzias latipes*) fish were bred and maintained as previously established (Loosli et al., 2000) (URL1). The animals used in the present study were from the inbred strain Cab. All experimental procedures were performed according to the guidelines of the German animal welfare law and approved by the local government (Tierschutzgesetz §11, Abs. 1, Nr. 1, husbandry permit number 35-9185.64/BH Wittbrodt).

### Plasmids

The variant DamL122A was created by site directed mutagenesis of the *E. coli Dam* gene using mutagenesis primers and flanking primers (Heckman and Pease, 2007) (Supplementary Table 1). The flexible linker was cloned as a dsOligo that encodes four GGGS amino acid repeats. The repeat sequences are flanked by Nhel and Spel sites. The mmGFP was amplified from plasmid pT2-*otp*ECR6_E1B::mmGFP (Gutierrez-Triana et al., 2014). All fragments were cloned into the pCS2+ vector (Rupp et al., 1994) as either amino or carboxy terminal fusions, followed by the SV40_polyadenylation signal of the pCS2 vector. Plasmid integrity was confirmed by sequencing.

We used the gene synthesis service of GeneArt (ThermoFisher Scientific) to obtain the optimized Dam sequence (oDam). In addition to codon optimization, cryptic splice sites were avoided (the DNA sequence is shown as an alignment to the unmodified DamL122A in Supplementary Fig.3). We replaced the DamL122A with the optimized *Dam* in the pCS2+ plasmids described above.
The DamL122A or oDam cassettes were excised using Agel and Notl from the pCS2+ plasmids and subcloned downstream of the 3.5 kb zebrafish *ubiquitin* promoter (Mosimann et al., 2011) in a Tol2_based plasmid (Kawakami et al., 1998) having *cmlc*.2::EGFP as the insertional reporter (Rembold et al., 2006).

### Microinjection

mRNA was synthesized using mMesage_mMachine SP6 kit (ThermoFisher Scientific, AM1340) on linearized pCS2+ templates. One cell stage zygotes were injected with mRNA at 10 ng/μl. The progenies of injected fish were maintained in ERM medium (Loosli et al., 2000) supplemented with penicillin_streptomycin (P0781, Sigma-Aldrich, 200 units/200 μg per ml of ERM). Embryos were collected at stage 22 (34 hpf at 28°C). Unfertilized and dead embryos were removed. Recombinant Ubiquitin promoter plasmids were injected into one-cell stage zygotes at 10 ng/μl in the presence of 10 ng/μl Tol2 transposase mRNA. The injected embryos were maintained in ERM medium supplemented with 0.2 mM N-phenylthiourea (Sigma-Aldrich, catalogue no. P7629) until hatching.

### Dpnl protection assay

pCS2+ plasmids carrying the cassettes coding for every particular *Dam* fusion protein were used to transform the *E. coli* strain C2925, deficient in the dam/dcm methylation system. As control, the pUC19 plasmid was used to transform C2925 cells and One Shot^®^ TOPIO cells, which have a normal methylation system. Bacterial genomic DNA was isolated from 3 ml LBamp cultures from individual colonies using the DNeasy Tissue kit (QIAGEN, Cat No. 69504). 1 μg of gDNA was digested with 10 units of DpnI (NEB, R0176S) for 1h at 37°C. The products were run in a 1% agarose gel.

### iDamlDseq protocol

*gDNA isolation*: Wash 20-30 embryos or tissue with lxERM or lxPBS respectively. Remove as much media as possible. Homogenize using pestle in 400 μl of TEN buffer (100 mM Tris-HCl pH8.5,10 mM EDTA, 200 mM NaCl, 1% SDS) plus 20 μl of 20mg/ml Proteinase_K. Incubate 0/N at 50-60°C. Cool down at RT for 5 min. Add 20 μl of 10 mg/ml RNase A (DNase and Proteinase-free, ThermoScientific, EN0531). Incubate 15 min at RT. Add 600 μl of phenol:chloroform:isoamylalcohol (25:24:1)(Roth, A156.1). Mix well by inversion. Incubate for 10 min at RT. Centrifuge at 10.000 rpm at RT for 20 min. Transfer the aqueous phase to a tube containing 600 μl of chloroform. Mix and centrifuge at 10.000 rpm for 10 min. Transfer the aqueous phase to a tube containing 600 μl of isopropanol. Mix and put at -20°C for 30 min. Centrifuge at 10.000 rpm at 4°C for 20 min. Remove supernatant and add 800 μl of cold 70% ethanol. Centrifuge at 14.000 rpm at RT for 10 min. Remove supernatant as much as possible and dry the pellet at 60°C for 10 min. Add 50 μl of prewarmed water (60°C). Incubate the tubes for 10-20 min at 60°C, gently flicking the tube sporadically until pellet is dissolved. Check OD260 and quality in a gel.

*DpnII Digestion and Alkaline phosphatase treatment*: In a 20 μl reaction, add 2 μl 10xNEB3.1 buffer, 1 μg of gDNA and 10 units of DpnII (NEB, R0543S). Incubate 6 hours at 37°C. Inactivate the enzyme at 65°C for 20min. To the inactive DpnII reaction add 23 μl H_2_O, 5 μl 10xAP buffer and 5 units of Antarctic phosphatase (NEB, M0289). Incubate reaction 1h at 37°C and inactivate at 70°C for 10 min. cleanup using column and elute in 12 μl of H_2_O.

*DpnI digestion*: In a 10 μl reaction, add 1 μl CutSmart buffer, 5 μl of DpnII/AP treated sample and 10 units of DpnI enzyme (NEB, R0176S). Exclude DpnI from the control tube. Incubate reaction at 37°C for 12 hours. Inactivate at 80°C for 20 min.

*Adaptor ligation*: To the DpnI +/– treated sample add 2μl lOx T4 ligase buffer, 1 μl of 50 μM dsOligos AdRt/b, 2.5 units of T4 DNA ligase (Thermo Scientific, EL0011) and 6.5 μl H_2_O. Incubate O/N at 16-18°C. Clean up using column and elute in 50 μl of H_2_O.

*LM-PCR*: In a 25 μl reaction, add 2.5 μl of lOx ThermoPol Buffer, 1 μl 10 mM dNTPS, 1 μl 10μM AdR_PCR primer (Supplementary Table 1), 5 μl of ligation sample and 1.25 units of Taq polymerase (NEB, M0267S). PCR program is as follows: 68°C 10 min, 1 cycle of 94°C 15 sec, 65°C 30 sec, 68°C 5 min and 20 to 30 cycles of 94°C 15 sec, 65°C 30 sec, 68°C 2 min. The number of cycles needs to be determined experimentally. Run 5 μl of PCR in a 1% agarose gel. A smear should appear only in the DpnI+ samples. Smear is typically from 200 bp to 2Kb.

*T7 exonuclease treatment*. To 30 μl of clean LM-PCR*, add 5 μl lOxCutSmart buffer, lOunits T7 exonuclease (NEB, M0263S) and 14 μl H_2_O. Incubate 1h at 25°C and cleanup using column in 20 μl of H_2_O. The sample is then ready for library preparation.

In order to obtain the minimum amount of DNA for deep sequencing it might be necessary to repeat the PCR step and pool the DNA samples.

### Sequencing

DNA samples were fragmented using the Covaris S2 sonicator in AFA microtubes. The library was prepared using the NEBNext Ultra DNA Library Prep kit for Illumina (E7370, NEB). Sequencing was performed in an Illumina HiSeq 2000 sequencing system.

### Data analysis

*Sequencing data processing*: Reads were mapped to the medaka genome (Kasahara et al., 2007) (oryLat2 assembly) using Bowtie2 (Langmead and Salzberg, 2012) with default parameters. Then the mapped reads were filtered with samtools (Li et al., 2009) to keep only those with a minimum mapping quality of 20.

*Identification of enriched regions*: First, the set of potential DpnI fragments was built from a BSgenome object for the oryLat2 assembly using the function vMatchPattern with the restriction site “GATC”. Only fragments spanning adjacent predicted restriction sites with length in the range from 200 to 2000 bases were considered. Next, the number of reads falling into each predicted fragment was counted for each one of the samples with the function summarizeOverlaps setting the parameter ignore.strand to TRUE. These counts were used to produce the correlation heatmap in Figure 1E. In order to discard fragments with spurious mapped reads, only fragments with a minimum number of reads relative to the fragment length were kept This threshold was computed as three times the total number of reads in all the fragments considered divided by the total length of them. After this selection, fragments that were not further apart than the smallest fragment length were joined together. With the resulting set of genomic regions, the read count was computed again with summarizeOverlaps and the resulting matrix was used to compute significant differences between samples using DESeq2 (Love et al., 2014). The R package iDEAR implements this data analysis pipeline and is available at https://bitbucket.org/iuanlmateo/idear.

*Motif enrichment* FIMO (Grant et al., 2011) was used to identified motif matches of the RAX binding motif (Jolma et al., 2013) (RAX_DBD) in the sequence within the coordinates of each identified region. The resulting regions were sorted by significance (from highest to lowest) and divided in 50 bins. This was done independently for each set of enriched or depleted Rx2 regions. For each bin, the ratio of regions with at least one match of the motif divided by the number of regions in the bin was computed. The shuffled sequences were generated by randomly permuting dinucleotides for each sequence extracted from the Rx2 regions.

### RT-PCR

10 hatchling larvae (10 dpf) per experimental group, expressing EGFP in the heart, were transferred to 1.5 ml tube and total RNA was isolated using Trizol reagent (ThermoFisher Scientific, 15596026) according to the manufacture's instructions. 1 (ig of DNase-treated total RNA was used to synthesize cDNA. PCR was done using Q5 polymerase (NEB, M0491) using 1xQ5 buffer, 200 μM dNTPs, 0.5μM RT_Ubi fwd primer, 0.5μM RT_mmGFP rev primer (Supplementary Table 1), 2 (il of cDNA and 0.02 u/μl Q5 polymerase. Cycling parameters were: 94°C 2min; 35 cycles, 94°C 15sec, 60°C 30 sec and 72°C 1min; 72°C 5 min. PCR products were analyzed in an agarose gel and sequenced.

## Acknowledgements

We thank Soojin Ryu for support during the initial DamID experiments. We highly appreciate the important input by Lazaro Centanin and Daigo Inoue, and the supporting feedback by the Wittbrodt lab.

### Author contributions

JAGT, JLM and JW designed the study. JAGT and DI performed the experiments. JLM developed iDEAR. JAGT, JLM and JW analyzed the data and wrote the manuscript.

### Data availability

The sequencing raw data are available at ArrayExpress with accession E-MTAB-4610.

